# Extracranial lymphatic–venous anastomosis remodels the choroid plexus–vascular axis in Alzheimer’s disease

**DOI:** 10.64898/2026.06.12.731621

**Authors:** Xinyu Pei, Subinuer Hujiabula, Zhengguo Zhang, Jiawang Zhang, Yupeng Hong, Ziguan Zhu, Renpeng Fang, Xiaodong Yang, Ye Lu, Liu Yang, Yingjun Zhao, Peng Guo

**Author notes:** E-mail: P. Guo; Y.J.Zhao L.Yang; Y. Lu.

## Abstract

Coordination of brain fluid transport, proteostasis and vascular function is essential for neural homeostasis, yet whether extracranial lymphatic outflow can actively shape central neurovascular states remains poorly understood. Here we identify a lymphatic–choroid plexus signaling axis through which extracranial drainage regulates intracranial proteostatic and vascular remodeling. Using cervical lymphatic–venous anastomosis (LVA) as a gain-of-function approach in an Alzheimer’s disease (AD) rat model, we show that enhancement of extracranial lymphatic outflow rapidly stabilizes cerebrospinal fluid–associated ventricular dynamics, selectively reduces oligomeric amyloid-β burden and improves locomotor performance. Single-cell transcriptomic profiling further reveals broad remodeling of the hippocampal neurovascular niche, marked by attenuation of inflammatory microglial programs and reinforcement of vascular communication networks. Mechanistically, LVA induces a secretory response in choroid plexus epithelial (CPE) cells characterized by co-upregulation of clusterin (CLU) and vascular endothelial growth factor (VEGF). This CLU–VEGF epithelial program links amyloid chaperoning to microvascular remodeling, accompanied by increased choroid plexus microvessel density and vascular caliber, preserved cerebral perfusion and restoration of neurovascular homeostasis. Together, these findings define the choroid plexus as a biomechanical–molecular transducer that connects extracranial lymphatic drainage with central proteostasis and vascular regulation. The lymphatic–choroid plexus axis provides a new framework for understanding how peripheral fluid drainage shapes brain homeostasis and suggests novel mechanism of action for treating neurodegenerative diseases.

## Introduction

Alzheimer’s disease (AD) is the leading cause of dementia and a major cause of disability in aging populations. More than 57 million people were living with dementia worldwide in 2021, and AD accounts for approximately 60–70% of cases, underscoring the scale of the clinical and societal challenge^1–3^. Recent anti-amyloid antibodies have established that modifying amyloid pathology can slow clinical progression in early AD, but the magnitude of benefit remains partial^4,5^. In the phase 3 CLARITY AD trial, lecanemab reduced clinical decline on the Clinical Dementia Rating–Sum of Boxes (CDR-SB) by 27% over 18 months, corresponding to an absolute difference of approximately 0.45 points. Similarly, donanemab slowed decline in early symptomatic AD, with a CDR-SB difference of approximately 0.67 points at 76 weeks in TRAILBLAZER-ALZ 2^6–8^. These advances are clinically meaningful, but they also highlight a central therapeutic gap: amyloid removal alone does not fully arrest neurodegeneration^9^, restore brain function, or correct the broader tissue-level disturbances that accompany AD^10,11^. Thus, there remains a need to explore new AD mechanisms that extend beyond direct amyloid targeting.

Recently, impaired brain fluid transport and neurovascular dysfunction have been identified as major contributors to AD pathophysiology^12^. The glymphatic pathway provides a route for cerebrospinal fluid (CSF) exchange with interstitial fluid and contributes to the clearance of soluble metabolites^13,14^, including amyloid-β. This process is closely linked to aquaporin-4-dependent perivascular water transport and becomes impaired with aging and neurodegenerative disease. The discovery of functional meningeal lymphatic vessels further revised the traditional view of the central nervous system as isolated from lymphatic drainage^15,16^ . Meningeal lymphatics drain CSF-associated solutes and immune cells toward deep cervical lymph nodes, and experimental impairment of this system aggravates age-associated and AD-like pathology^17–19^.In parallel, AD is associated with substantial neurovascular dysfunction. Cerebral blood flow reductions of approximately 20–30% have been reported in AD, and perfusion deficits can appear early in the disease process and correlate with cognitive impairment. These observations suggest that fluid clearance, lymphatic drainage and vascular perfusion are not independent features of AD, but interconnected components of a vulnerable brain homeostatic system.

Cervical LVA offers a surgical means to redirect extracranial lymphatic outflow into the venous circulation^20^. Although lymphatic–venous bypass is an established strategy for decompression of peripheral lymphatic disorders, its application to AD remains highly controversial. In July 2025, the National Health Commission of China prohibited the clinical use of deep cervical lymphatic vessel/node–venous anastomosis for AD treatment, citing its early exploratory status, unclear indications and contraindications, and insufficient high-quality evidence for safety and efficacy^21^. These concerns highlight a central knowledge gap: whether extracranial lymphatic redirection can reproducibly engage disease-relevant brain biology.

Here, we address this challenge by using cervical LVA as a gain-of-function strategy to determine whether extracranial lymphatic redirection can engage disease-relevant brain mechanisms in an AD rat model. By integrating microsurgery, longitudinal magnetic resonance imaging (MRI), Aβ species-specific pathology, gait analysis, and single-cell profiling analyses, we show that LVA remodels the AD brain microenvironment within 14 days. LVA stabilized CSF-associated ventricular structures, preferentially reduced oligomeric Aβ, improved gait performance and preserved cerebral perfusion. Mechanistically, LVA activated a CLU–VEGF epithelial program in the choroid plexus, linking fluid-dynamic cues to neurovascular remodeling and amyloid handling^22,23^. These findings identify a lymphatic–choroid plexus–vascular axis by which extracranial lymphatic outflow can regulate central proteostatic and neurovascular homeostasis in AD.

## Results

### Cervical LVA establishes a functional extracranial drainage pathway and improves gait performance in AD rats

To evaluate the therapeutic benefits of cervical LVA for AD, we established a unilateral cervical LVA model in 2-month-old AppNL-G-F knock-in AD rats^24^ , a disease stage characterized by early Aβ pathology and emerging neurovascular dysfunction^25,26^. Animals were randomly assigned to LVA or sham-operated control groups and subjected to a longitudinal experimental workflow integrating baseline assessment, serial plasma biomarker profiling, structural and perfusion MRI, behavioral testing, histological analysis and single-cell transcriptomic profiling (Fig. 1A). This design allowed us to evaluate both the technical feasibility of cervical LVA and its early functional and biological consequences within a defined 14-day post-surgical window^27^.

**Figure 1.**
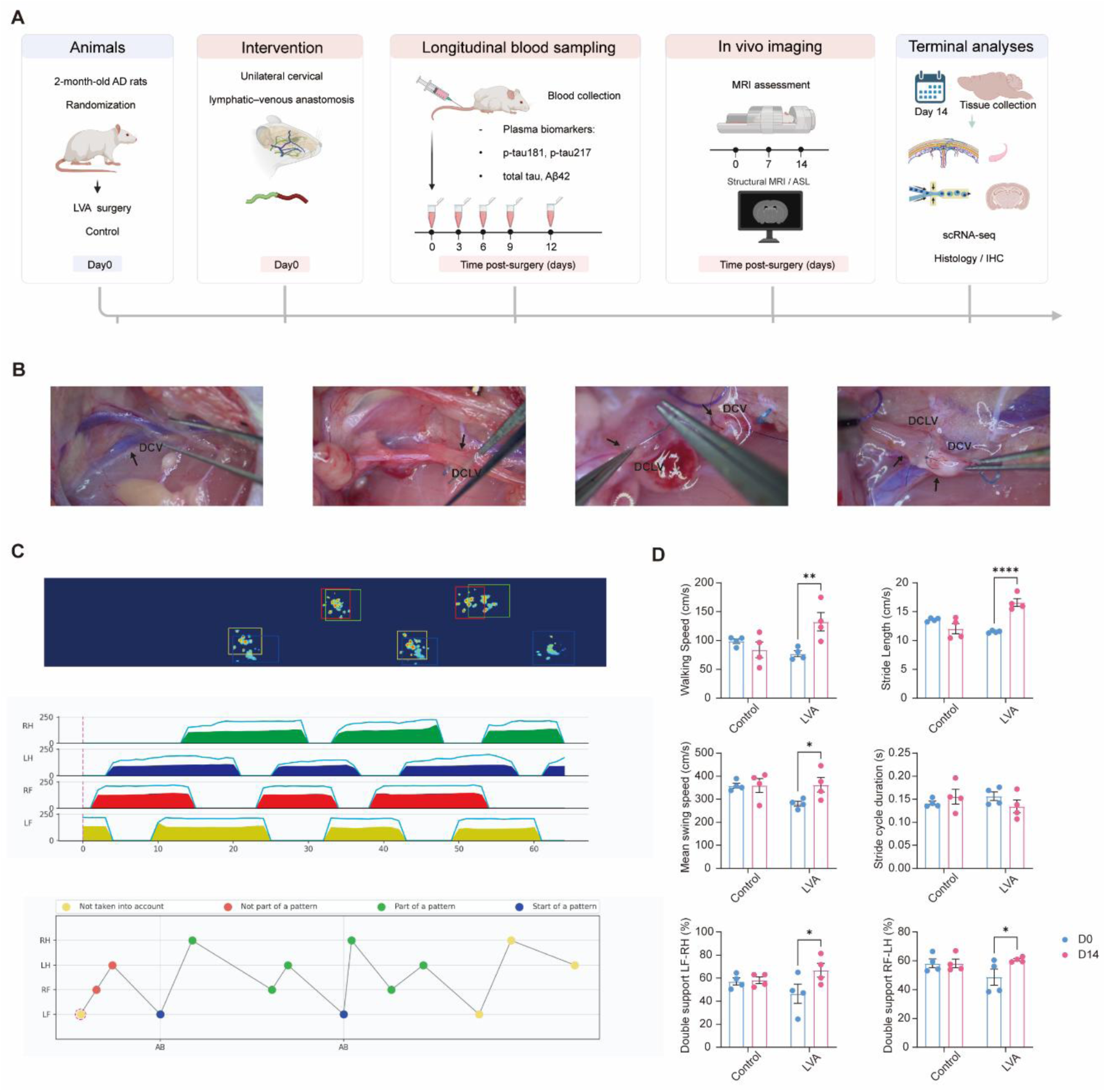
LVA experimental design, surgical procedure, and gait assessment. (A) Experimental workflow showing baseline assessment, unilateral cervical LVA or sham surgery, serial blood collection, longitudinal MRI, gait testing and terminal tissue analyses. (B) Representative intraoperative images showing exposure of the DCV and DCLV, followed by construction of an end-to-end lymphatic–venous anastomosis. (C) Representative gait footprint and temporal paw-placement traces at day 14 after surgery. (D) Quantification of gait parameters at baseline and day 14, including walking speed, stride length, mean swing speed, stride cycle duration and diagonal double-support phase (LF–RH and RF–LH). Data are presented as mean±SEM (n = 4 per group). Statistical significance is indicated in the panels.

Under high-magnification microsurgery, the deep cervical vein (DCV) was exposed and isolated, and the adjacent deep cervical lymphatic vessel (DCLV) was identified by its translucent wall, small caliber and low intraluminal pressure. An end-to-end anastomosis was then constructed between the DCLV and DCV using 12–0 nylon sutures with minimal tissue manipulation (Fig. 1B). Intraoperative inspection showed immediate and continuous lymphatic-to-venous flow without leakage, confirming the establishment of a patent lymphatic–venous junction. All LVA procedures were technically successful, whereas sham-operated rats underwent identical cervical exposure without vessel transection or anastomosis. Animals in both groups tolerated the procedure well, with no overt post-operative morbidity during the observation period. These results demonstrate that cervical LVA provides a reliable gain-of-function approach to divert deep cervical lymphatic outflow into the venous circulation in AD rats.

We next asked whether this surgically established extracranial drainage pathway was associated with early functional improvement. Gait performance was assessed before surgery and 14 days after intervention using an automated gait analysis system (Fig. 1C). At baseline, LVA and sham groups showed comparable locomotor parameters, indicating similar pre-operative motor function. By day 14, LVA-treated rats exhibited improved gait performance compared with sham-operated controls, with walking speed increased by approximately 60–70% and diagonal double-support phases increased by approximately 25–35% relative to baseline, accompanied by longer stride length, higher swing speed and shorter stride cycle duration (Fig. 1D). These changes indicate a more efficient and dynamically stable gait pattern after LVA.

### Cervical LVA stabilizes CSF-associated brain structures and reduces oligomeric Aβ **burden**

We next examined whether enhanced extracranial lymphatic outflow by cervical LVA could influence intracranial fluid homeostasis and amyloid pathology^18^. Longitudinal 7.0-T T2-weighted MRI was performed before surgery and at days 7 and 14 after intervention to monitor CSF-associated brain structures over time (Fig. 2A,B). Because baseline ventricular size varied between animals and groups, ventricular and hippocampal measurements were normalized to the corresponding D0 value of each animal and expressed as relative percentage change from baseline (Fig. 2C). Sham-operated AD rats showed progressive enlargement of the lateral ventricles and third ventricle, accompanied by a relative reduction in hippocampal area over time. In contrast, LVA-treated rats showed markedly attenuated ventricular expansion and better preservation of hippocampal morphology, with the strongest stabilization observed in CSF-associated ventricular measures at day 7 and maintained through day 14. These data indicate that cervical LVA stabilizes CSF-associated structural dynamics.

**Figure 2.**
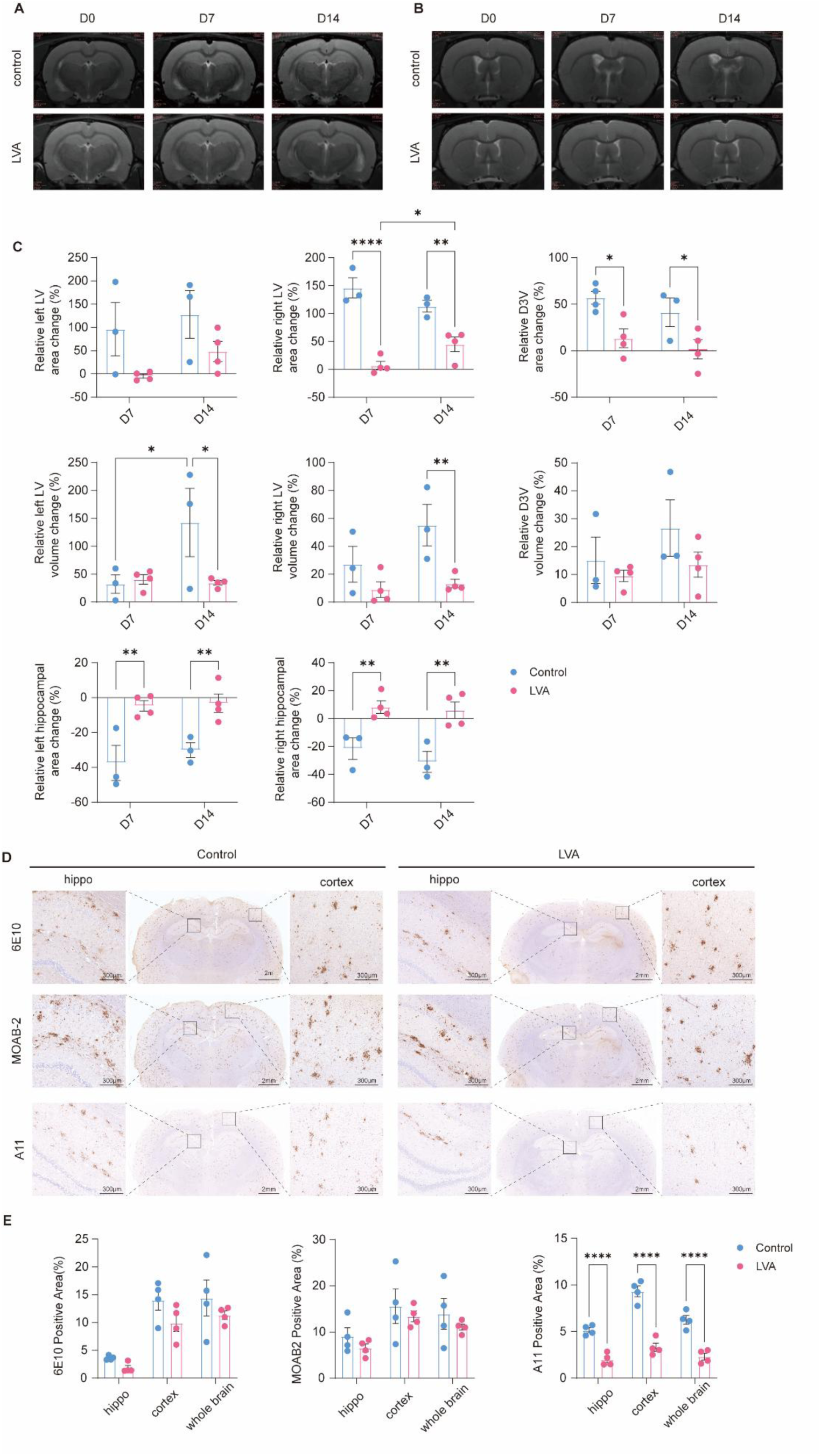
Cervical LVA stabilizes CSF-associated brain structures and reduces cerebral amyloid burden. (A,B) Representative longitudinal T2-weighted coronal MRI images acquired at baseline (D0), day 7 (D7) and day 14 (D14) after sham or LVA surgery, showing anatomically matched sections through the lateral ventricles (LV) and third ventricle (3V). (C) MRI-derived hippocampal and ventricular measurements normalized to the corresponding D0 value of each animal and expressed as relative percentage change from baseline. LVA attenuated progressive LV and 3V enlargement and preserved hippocampal morphology compared with sham controls. (D) Representative immunohistochemistry of coronal brain sections at D14 showing Aβ-related pathology in the hippocampus and cortex, detected by 6E10 (total Aβ/APP-related signal), MOAB-2 (deposited/fibrillar Aβ) and A11 (oligomeric Aβ). Scale bars, 2 mm (whole brain) and 300 μm (insets). (E) Quantification of 6E10-, MOAB-2- and A11-positive area in the hippocampus, cortex and whole brain, showing preferential reduction of oligomeric Aβ after LVA. Data are presented as mean±SEM. Statistical analysis and significance are indicated in the panels. Statistical analysis by two-way ANOVA.

We next assessed whether these changes were accompanied by altered amyloid burden. Immunohistochemical (IHC) staining was performed using antibodies recognizing distinct Aβ-related species, including 6E10 for total Aβ/APP-related signal^28^, MOAB-2 for Aβ deposits^29^, and A11 for oligomeric Aβ (Fig. 2D). LVA produced only modest reductions in 6E10- and MOAB-2-positive Aβ signals, suggesting that total or deposited Aβ was not globally eliminated within two weeks. In contrast, A11-positive oligomeric Aβ was markedly reduced after LVA across the hippocampus, cortex and whole-brain sections, with highly significant differences compared with sham controls (****P < 0.0001; Fig. 2E). This pattern suggests that LVA preferentially affects soluble or oligomeric Aβ species rather than reducing all amyloid-related signals indiscriminately.

### Cervical LVA induces coordinated neurovascular remodeling in the hippocampus

To delineate the mechanism underlying LVA-based neuroprotection, we integrated single-cell transcriptomics, vascular histology and longitudinal perfusion MRI, with a focus on the hippocampus and CSF-associated compartments. Single-cell RNA sequencing of 73,140 hippocampal cells resolved the major neural, glial, vascular and immune populations, including microglia, oligodendrocytes, OPCs, astrocytes, neurons, immature neurons, endothelial cells, mural cells, CPE cells, border-associated macrophages and neutrophils (Fig. 3A,B). Comparative analysis revealed that LVA did not act on a single cell lineage, but induced a broad shift in the hippocampal cellular ecosystem, with changes in the relative abundance of immune, glial and vascular-associated populations (Fig. 3C).

**Figure 3.**
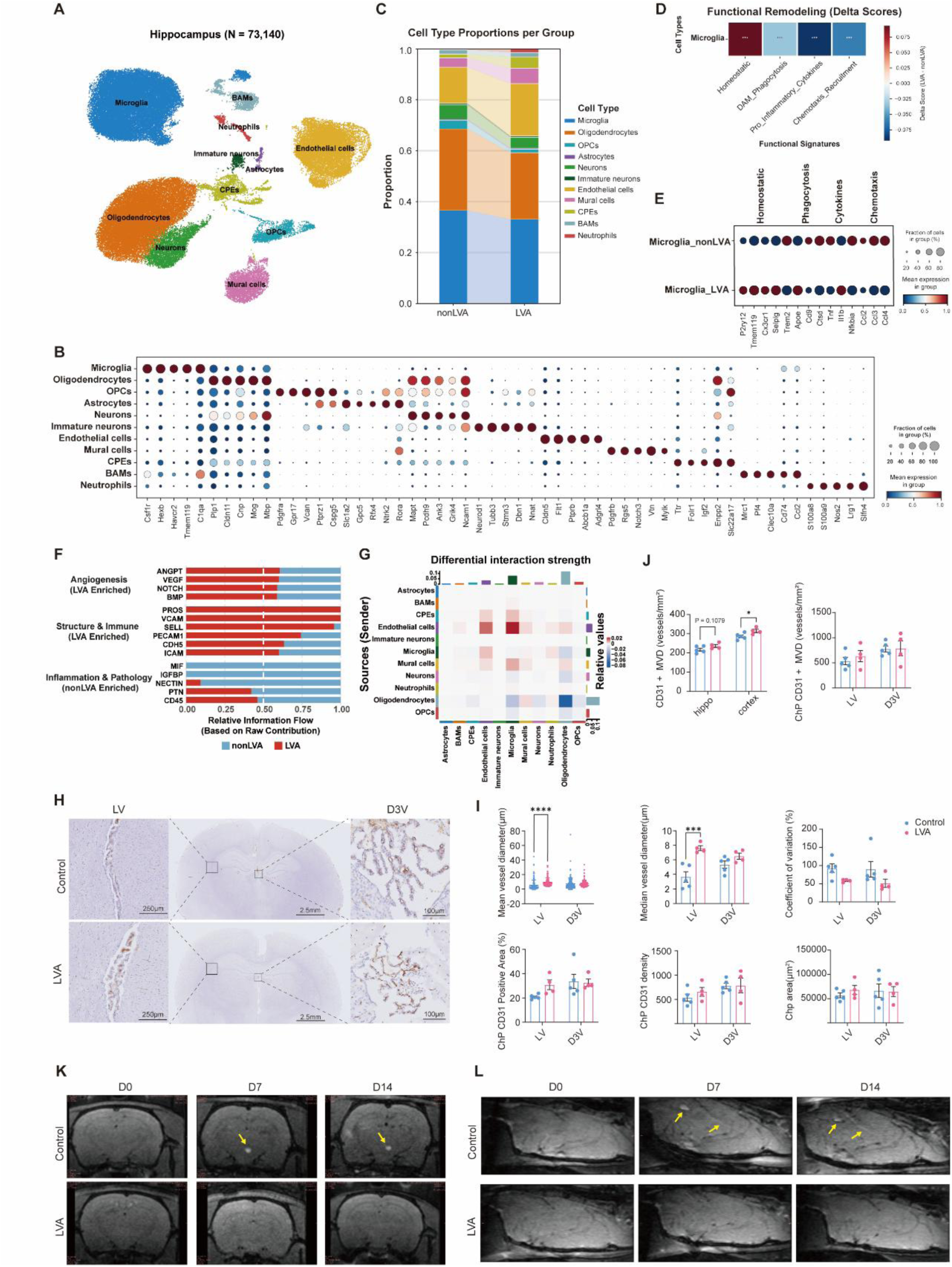
Cervical LVA induces coordinated neurovascular remodeling and stabilizes cerebral perfusion in the hippocampus. (A) UMAP projection of 73,140 hippocampal single-cell transcriptomes, resolving major neural, glial, vascular and immune cell populations. (B) Dot plot of canonical marker genes used to annotate the identified hippocampal cell populations. (C) Bar plot showing the relative proportions of major hippocampal cell populations in control and LVA-treated rats. (D,E) Functional scoring (D) and marker-gene dot plot (E) showing microglial reprogramming after LVA, characterized by reduced inflammatory/chemotactic signatures and relative restoration of homeostatic markers. (F,G) Relative information flow (F) and differential interaction-strength heatmap (G) showing reorganization of intercellular communication, with enrichment of vascular signaling programs after LVA. (H) Representative CD31-stained coronal brain sections showing CD31+ vasculature in the lateral ventricle choroid plexus (LV-ChP) and dorsal third ventricle choroid plexus (D3V-ChP). Scale bars as indicated. (I) Quantification of ChP vascular morphometric parameters, including mean vessel diameter, median vessel diameter, coefficient of variation of vessel diameter, CD31-positive area, ChP CD31 density and total ChP area. LVA markedly increased LV-ChP vessel diameter and showed an upward trend in CD31+ vascular density without gross expansion of ChP area. (J) Quantification of CD31+ microvessel density in the hippocampus, cortex, LV-ChP and D3V-ChP. LVA significantly increased cortical MVD and induced an upward trend of MVD in hippocampal and ChP regions. (K) Representative longitudinal ASL-MRI maps from D0 to D14; yellow arrows indicate focal perfusion abnormalities in controls. (L) pCASL imaging showing attenuation of regional perfusion abnormalities and maintenance of hemodynamic stability in LVA-treated rats. Data are presented as mean ± SEM. Statistical significance was determined by two-way ANOVA.

Given the central role of neuroinflammation in AD pathology, we first examined microglial transcriptional states. Functional scoring analysis revealed a marked suppression of pro-inflammatory programs following LVA, including cytokine production and chemotactic recruitment pathways, accompanied by a relative increase in homeostatic signatures (Fig. 3D). Consistently, expression of canonical homeostatic microglial markers (e.g., P2ry12, Tmem119) was elevated^30^, whereas pro-inflammatory genes such as Il1b and Ccl3 were reduced compared with controls ^31^(Fig. 3E). These findings indicate that LVA alleviates neuroinflammation by shifting microglia from a disease-associated^18^, pro-inflammatory state toward a homeostatic phenotype.

We next asked whether this microglial remodeling was accompanied by altered intercellular communication. Ligand–receptor analysis showed a redistribution of signaling pathways after LVA (Fig. 3F). Angiogenesis- and vascular-structure-related pathways, including VEGF- and NOTCH-associated signaling, were enriched in the LVA group, whereas inflammatory and pathology-associated pathways were relatively enriched in controls. Differential interaction analysis further indicated strengthened communication among endothelial cells, CPEs and other hippocampal niche components (Fig. 3G), supporting enhanced neurovascular crosstalk after lymphatic redirection and positioning the choroid plexus–vascular interface as a central signaling hub downstream of LVA.

We examined whether the VEGF-centered vascular signaling response was accompanied by CD31⁺ vascular remodeling. CD31 IHC revealed marked vascular changes in the choroid plexus, most prominently in the LV-ChP compartment (Fig. 3H). LVA significantly increased LV-ChP vessel diameter, with mean and median diameters rising by 75.0 % and 108.2 %, respectively (Fig. 3I). Vessel diameter in the D3V-ChP showed a similar but weaker upward trend, while the coefficient of variation decreased by 37.2%, indicating a more uniform vascular architecture. In addition to caliber expansion, CD31⁺ microvessel density (MVD) was significantly increased in the cortex by 9.8%, with upward trends in the hippocampus and ChP, especially LV-ChP, whereas total ChP area remained unchanged (Fig. 3J). These findings indicate that cervical LVA drives region-dependent vascular remodeling, dominated by significant LV-ChP vessel enlargement and accompanied by selective microvascular expansion.

Finally, we evaluated whether this vascular remodeling was associated with functional preservation of cerebral perfusion. Longitudinal ASL-MRI showed that control AD rats developed increasingly heterogeneous perfusion patterns over time, with focal hypoperfused regions appearing at days 7 and 14 (Fig. 3K). In contrast, LVA-treated rats maintained a more homogeneous perfusion pattern throughout the 14-day observation period. Consistent with this finding, pCASL imaging showed attenuation of regional perfusion deficits after LVA, whereas control rats displayed persistent focal perfusion abnormalities (Fig. 3L).

Together, these results show that cervical LVA induces coordinated remodeling of the hippocampal neurovascular niche^32^. This remodeling is characterized by partial restoration of microglial homeostasis, amplification of VEGF-centered vascular communication, normalization of brain microvascular structure and preservation of cerebral perfusion. Thus, extracranial lymphatic–venous redirection appears to influence AD pathology not only through fluid drainage, but also by engaging neurovascular repair programs within the hippocampal–choroid plexus axis.

### Cervical LVA activates a mechanosensitive choroid plexus–vascular signaling axis

To further define the molecular pathway linking extracranial lymphatic drainage to neurovascular remodeling, we focused on CPEs and endothelial cells^33^, two cell populations positioned at the interface of CSF homeostasis, amyloid handling and vascular regulation. Single-cell transcriptomic analysis revealed that LVA broadly reprogrammed CPE function toward a clearance-competent and neurotrophic secretory state (Fig. 4A). Genes involved in amyloid-associated binding and trafficking molecules, including Lrp2, Apoe ^34,35^ and Clu^36,37^, were upregulated, indicating reinforcement of Aβ-handling pathways. LVA also increased trophic and vascular regulatory factors such as Vegfa and Igf2^38^, suggesting that CPEs acquire a neurovascular-supportive secretory phenotype after lymphatic–venous redirection.

**Figure 4.**
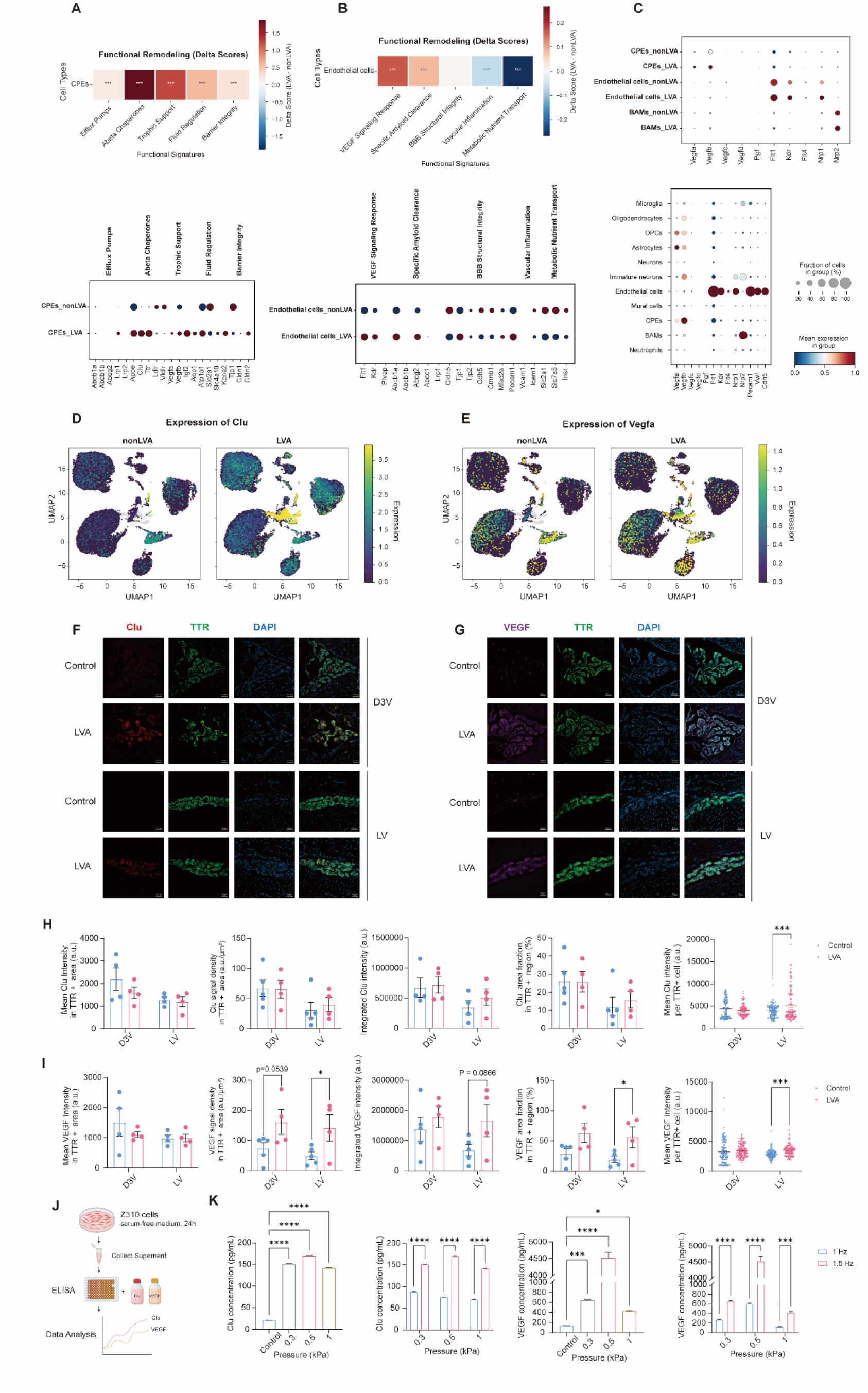
Cervical LVA reprograms choroid plexus and endothelial functions via VEGF–CLU signaling and biomechanical feedback. (A) Heatmap of GSVA enrichment scores in choroid plexus epithelial cells (CPEs), showing upregulated pathways related to efflux transporters, Aβ chaperones, and trophic factor secretion post-LVA. (B) Functional enrichment analysis of endothelial cells, illustrating enhanced expression of tight junction components and metabolic transport pathways following LVA. (C) Dot plot illustrating ligand–receptor interactions, highlighting the activation of the VEGF signaling axis (Vegfa–Flt1/Kdr) between CPEs and endothelial cells. (D–E) UMAP plots showing the expression and cluster expansion of Clu (D) and Vegfa (E) in non-LVA versus LVA samples. (F–G) Representative immunofluorescence images of the choroid plexus in the lateral ventricle (LV) and third ventricle (D3V). Tissues were stained for TTR (green), CLU (red), and VEGF (magenta), showing LVA-induced protein upregulation in TTR⁺ epithelial cells. Scale bars =50μm. (H–I) Quantitative analysis of CLU intensity per TTR⁺ cell (H) and VEGF signal density (I), confirming significant increases in the LVA group. (J) Experimental schematic of the in vitro biomechanical stimulation model using Z310 cells subjected to controlled pressure amplitudes and pulsatile frequencies. (K) ELISA quantification of secreted CLU and VEGF under varying mechanical stimuli, showing optimal secretion at 0.3–0.5 kPa and 1.5 Hz. For in vivo immunofluorescence quantification in H–I, data are presented as mean ± SEM. For in vitro ELISA experiments in K, data are presented as mean ± SD. Statistical significance was determined by two -way ANOVA (*p < 0.05, **p < 0.01, ***p < 0.001, ****p < 0.0001).

Endothelial cells showed a coordinated phenotype shift consistent with improved VEGF-signaling response and specific amyloid-clearance, while the BBB structural-integrity signature remained largely unchanged (Fig. 4B). LVA increased the expression of angiogenesis-related genes, including Flt1 and Kdr together with specific amyloid-clearance genes Abcb1a and Abcg2. By contrast, vascular-inflammation and metabolic nutrient-transport scores decreased. Thus, the endothelial response to LVA appears to reflect a restricted VEGF-linked and amyloid-clearance-associated remodeling program, rather than a generalized strengthening of barrier structure or nutrient-exchange function.

We next asked which intercellular signals might couple CPE activation to endothelial remodeling. Ligand–receptor analysis identified VEGF signaling as a prominent LVA-responsive pathway within the choroid plexus–vascular niche (Fig. 4C). Increased Vegfa expression in CPEs, together with increased endothelial expression of VEGF receptors Flt1 and Kdr^33^, supported enhanced epithelial-to-endothelial paracrine communication. Notably, Clu was predominantly enriched in CPEs and further increased after LVA. UMAP visualization further confirmed that LVA increased Clu expression within CPE-enriched epithelial states, while Vegfa expression was also enhanced in the choroid plexus cellular compartment (Fig. 4D,E). Notably, UMAP feature plots showed enhanced Clu expression in CPE-enriched epithelial states after LVA, accompanied by increased Vegfa expression across the choroid plexus cellular compartment (Fig. 4D,E). We validated these transcriptional findings at the protein level by immunofluorescence staining of choroid plexus tissues. In both the LV and D3V regions, LVA markedly increased CLU and VEGF signals within TTR-positive epithelial cells (Fig. 4F,G). Quantification showed an approximately 2–3-fold increase in CLU intensity per TTR-positive cell and an approximately two-fold increase in VEGF signal density after LVA (Fig. 4H,I). These signals were enriched along epithelial folds, consistent with activation of choroid plexus secretory domains. Together, these data indicate that cervical LVA drives CPEs toward an activated CLU–VEGF-producing epithelial state.

Finally, we tested whether altered fluid-dynamic forces could directly induce this CPE secretory program. Using a DZ-21 TSS organ/tissue culture system, rat CPE Z310 cells were exposed to defined cyclic hydrostatic pressure amplitudes and frequencies (Fig. 4J). Low-amplitude, CSF-like cyclic pressure stimulation at 0.3–0.5 kPa, corresponding to approximately 2.25–3.75 mmHg, markedly increased CLU secretion by approximately 7–8-fold and VEGF secretion by up to approximately 45-fold compared with static controls, whereas this response was attenuated at 1.0 kPa (Fig. 4K). Increasing the pulsatile frequency from 1.0 to 1.5 Hz further enhanced both CLU and VEGF secretion, suggesting that CPEs are sensitive not only to pressure amplitude but also to pulsatile frequency within a range relevant to cardiac- and vascular-driven CSF pulsatility. Thus, CPEs are intrinsically capable of sensing low-amplitude cyclic pressure cues and converting them into a CLU–VEGF secretory response, providing a plausible mechanistic link between altered extracranial drainage and choroid plexus–vascular remodeling after LVA.

### Peripheral biomarkers reflect systemic responses following cervical LVA

To determine whether the effects of cervical LVA could be detected by minimally invasive systemic readouts, we longitudinally quantified plasma Aβ42 and VEGF, with values normalized to each animal’s pre-surgical baseline (Fig. 5A,B). In sham-operated controls, plasma Aβ42 remained largely stable or declined over the post-surgical period. In contrast, LVA-treated AD rats showed a sustained increase in plasma Aβ42 relative to baseline, with most individual trajectories remaining above control levels after surgery (Fig. 5A). This pattern is consistent with increased peripheral mobilization or redistribution of soluble amyloid species following extracranial lymphatic–venous redirection.

**Figure 5.**
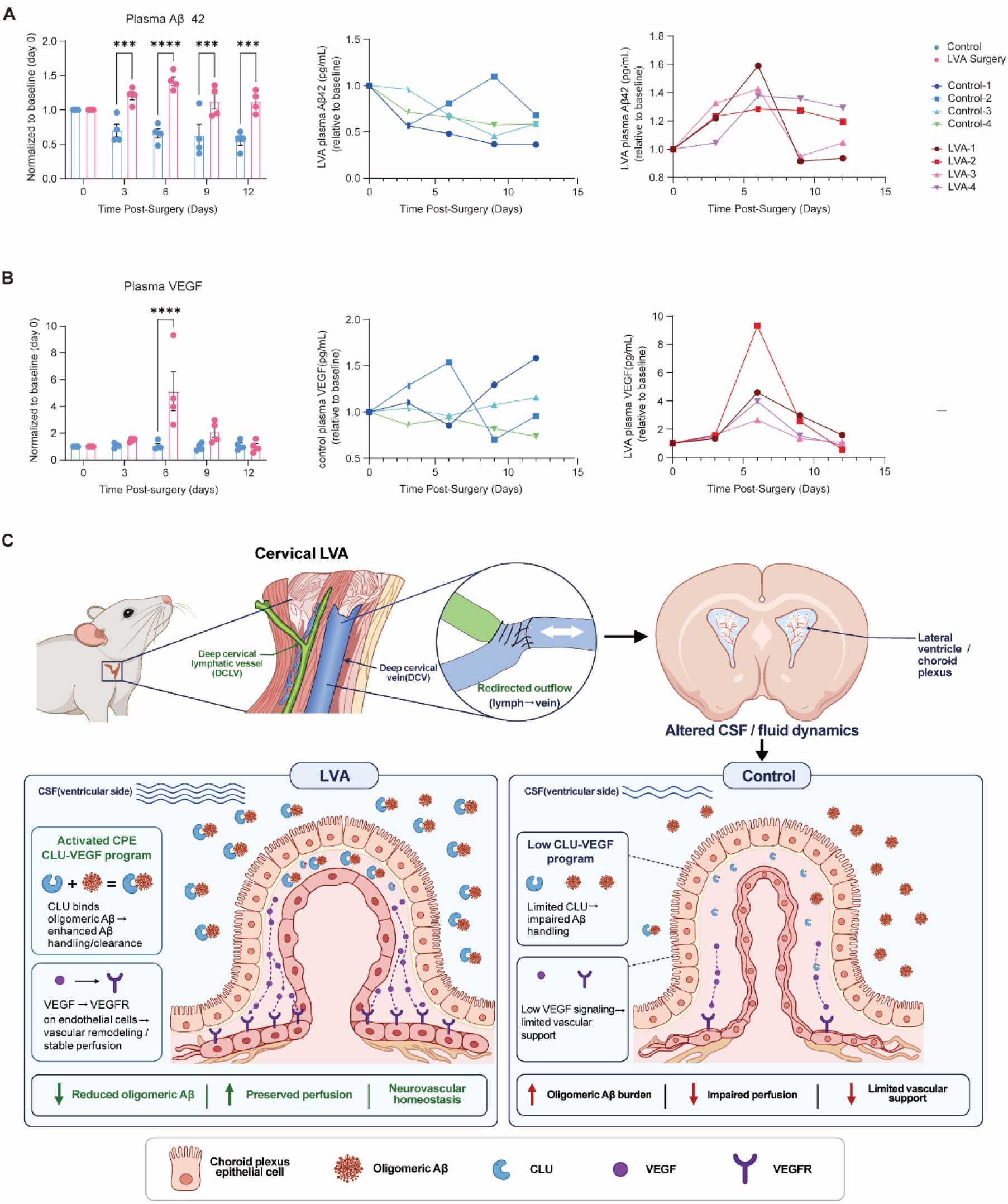
Peripheral biomarkers reflect systemic neuroprotective effects and validate LVA efficacy. (A) Longitudinal plasma Aβ42 levels normalized to each animal’s baseline (D0), shown as group summaries and individual trajectories in sham and LVA-treated AD rats at days 3, 6, 9 and 12 after surgery. LVA increased plasma Aβ42 relative to controls, consistent with enhanced mobilization of soluble amyloid species. (B) Longitudinal plasma VEGF levels normalized to baseline, shown as group summaries and individual trajectories. LVA induced a transient VEGF increase during the early post-operative period, consistent with activation of vascular signaling. (C) Schematic model of the lymphatic–choroid plexus–vascular axis. Cervical LVA redirects extracranial lymphatic outflow, alters CSF/fluid dynamics and activates a mechanosensitive CLU–VEGF program in choroid plexus epithelial cells, leading to improved amyloid handling, vascular remodeling, preserved perfusion and systemic biomarker changes detectable in peripheral blood.

Plasma VEGF showed a distinct temporal profile. LVA induced a transient increase in circulating VEGF, with the strongest elevation occurring during the early post-operative period followed by a gradual return toward baseline (Fig. 5B). This peripheral VEGF response was concordant with the CPE-derived VEGF activation observed by transcriptomic and tissue levels (Figs. 3-4). These findings extend the CLU–VEGF axis from a local choroid plexus response to a systemically detectable signature, indicating that the mechanosensitive epithelial program triggered by lymphatic redirection is reflected in the peripheral circulation.

## Discussion

AD is increasingly recognized not only as a disorder of amyloid deposition and neuronal degeneration, but also as a condition shaped by impaired brain fluid transport, neurovascular dysfunction and chronic neuroimmune remodeling^16,39^. Previous work on glymphatic and meningeal lymphatic systems has shown that CSF–interstitial fluid exchange contributes to the clearance of soluble metabolites and protein aggregates from the brain^18,40^. AQP4-dependent glymphatic transport provides a well-established example, as AQP4 deficiency reduces interstitial solute clearance by approximately 70%, indicating that fluid transport is an active determinant of brain waste handling rather than a secondary consequence of pathology^40,41^ . Consistent with this concept, our MRI data showed that LVA attenuated ventricular expansion, most clearly in the D3V at day 7, while hippocampal area remained largely unchanged. Histologically, LVA had only modest effects on total or deposited Aβ detected by 6E10 and MOAB-2, but markedly reduced A11-positive oligomeric Aβ across hippocampal, cortical and whole-brain regions. Thus, LVA does not appear to acutely remove established amyloid deposits within two weeks; instead, it preferentially affects more dynamic and fluid-sensitive Aβ species, particularly oligomeric Aβ, which is closely associated with early synaptic toxicity. The accompanying gait improvement, including increased walking speed and diagonal double-support phases, further suggests that lymphatic redirection can produce measurable neurological benefits within an early disease window.

A key finding of this study is that LVA acts beyond passive lymphatic drainage^42–44^. Single-cell profiling of hippocampal cells revealed broad remodeling of immune, glial and vascular-associated populations after LVA. The strongest evidence for neurovascular modulation, however, came from vascular imaging and histology. CD31 staining showed a significantly increase in median microvascular diameter and MVD in the LV-ChP region. This pattern is more consistent with remodeling or normalization of pre-existing choroid plexus vessels than with broad de novo angiogenesis^45^. Functionally, this vascular remodeling was accompanied by more homogeneous ASL and pCASL perfusion patterns, whereas sham-operated AD rats developed focal hypoperfused regions over time^46^. These data link extracranial lymphatic redirection to stabilization of cerebral hemodynamics.

The choroid plexus was identified as a key intermediary between altered lymphatic outflow and central neurovascular remodeling. This interpretation is supported by the anatomical position of the choroid plexus at the interface of CSF production, blood–CSF barrier regulation and soluble factor secretion^47,48^. In our study, immunofluorescence analysis showed that LVA markedly increased CLU and VEGF signals within TTR-positive CPE cells in both the lateral ventricle and dorsal third ventricle regions. Quantitatively, CLU intensity per TTR-positive cell increased by approximately 2–3-fold, whereas VEGF signal density increased by approximately twofold. CLU provides a plausible link to Aβ handling, whereas VEGF provides a direct link to vascular remodeling^49,50^. Therefore, the CLU–VEGF epithelial response offers a coherent mechanism by which the choroid plexus may couple improved protein handling to neurovascular repair.

CLU and VEGF signaling emerged as candidate molecular pathways linking extracranial lymphatic redirection to amyloid handling and vascular remodeling^51^ . After LVA, CLU and VEGF were co-induced in TTR-positive choroid plexus epithelial cells, with CLU intensity increasing by approximately 2–3-fold and VEGF signal density by approximately twofold. This coordinated response is biologically plausible: CLU has been implicated in Aβ binding, transport and clearance, whereas VEGF can regulate endothelial remodeling and vascular adaptation. Notably, VEGF induction was accompanied by increased vessel caliber, MVD and improved perfusion homogeneity, suggesting a spatially restricted reparative response rather than uncontrolled angiogenesis.

The pressure-stimulation experiments further suggest that this epithelial program may be mechanically tuned^52^. In Z310 cells, moderate cyclic hydrostatic pressure induced CLU and VEGF secretion, whereas higher pressure attenuated the response; increasing pulsatile frequency from 1.0 to 1.5 Hz further enhanced secretion. These findings indicate that CPEs can translate fluid-dynamic cues into a secretory output that resembles the in vivo CLU–VEGF response after LVA. Thus, cervical LVA may act not only by redirecting lymphatic drainage, but also by reshaping the mechanical environment sensed by the choroid plexus, thereby coupling Aβ handling to neurovascular repair. Future studies will be required to identify the upstream mechanosensors and to determine whether CLU or VEGF signaling is necessary for the protective effects of LVA.

Several limitations should be noted. First, the study was performed in an early-stage AD rat model over a short 14-day interval; whether the observed effects persist or translate into long-term cognitive benefit remains unknown. Second, although the data implicate a choroid plexus CLU–VEGF response, direct causal experiments are still needed, including blockade of VEGF signaling, CLU perturbation or selective manipulation of choroid plexus mechanotransduction. Third, gait improvement is an objective functional readout, but additional cognitive assays will be required to establish effects on learning and memory. Fourth, the vascular data support structural remodeling and perfusion stabilization, but do not yet define BBB permeability, vessel maturity or ultrastructural integrity. Finally, species differences in cervical lymphatic anatomy, CSF dynamics and choroid plexus biology must be considered before extrapolating these findings to human AD.

In summary, this study identifies a lymphatic–choroid plexus–vascular axis through which cervical LVA remodels the AD brain microenvironment. Rather than acting solely as a lymphatic drainage procedure, LVA appears to engage a mechanosensitive CPE program marked by CLU and VEGF activation, leading to reduced oligomeric Aβ, choroid plexus vascular remodeling and preservation of cerebral perfusion. These findings shift the conceptual framework of lymphatic intervention in AD from waste clearance alone to neurovascular niche remodeling and provide a basis for future biomarker-guided evaluation of extracranial lymphatic modulation in neurodegenerative disease.

## Materials and Methods

### Animals

All experimental procedures were approved by the Institutional Animal Care and Use Committee of Hangzhou Institute of Medicine, Chinese Academy of Sciences (protocol number: AP2025-06-0522),and were conducted in accordance with national and institutional guidelines for the care and use of laboratory animals. In this study, male App^NL-G-F^ knock-in rats were used as an Alzheimer’s disease (AD) model^24^. This line was generated by CRISPR/Cas9-mediated replacement of the endogenous rat *App* locus with a humanized *APP* sequence harboring three familial AD-associated mutations: Swedish (KM670/671NL), Arctic (E693G), and Beyreuther/Iberian (I716F). Expression of the humanized *APP* allele was driven by the endogenous rat *App* promoter, thereby preserving physiological expression levels and spatial distribution patterns^24^. All animals were obtained from LFP Biosciences (Beijing, China). Rats were 2 months of age at the beginning of the experiments, corresponding to an early pathological stage in this model, characterized by the onset of amyloid-β (Aβ) oligomerization and initial plaque deposition. Animals were housed under specific pathogen–free conditions on a 12-h light/dark cycle and had ad libitum access to food and water.

### Cervical deep lymphatic–venous anastomosis (LVA)

Unilateral cervical deep lymphatic–venous anastomosis (LVA) was performed in 2-month-old AD rats^20^. Animals were anesthetized with isoflurane and maintained on a thermostatically controlled heating pad throughout the procedure. After removal of cervical hair and sterilization, a midline cervical incision was made from the submandibular region. Under an operating microscope, the deep cervical vein and the adjacent deep cervical lymphatic vessel were microsurgically exposed. Temporary ligatures were applied to the proximal and distal segments of the deep cervical vein relative to its junction with the common jugular vein to achieve transient vascular control. Permanent ligatures were placed at both ends of the deep cervical lymphatic vessel. The vein and lymphatic vessel were then transected at the ligated sites, and the lumens were gently irrigated with heparinized saline (125 U/mL). An end-to-end lymphatic–venous anastomosis was constructed using 3–4 interrupted microsutures. Following completion of the anastomosis, venous patency was confirmed and the temporary venous ligatures were released to restore blood flow. The surgical field was disinfected, the incision was closed in layers, and animals were monitored until full recovery from anesthesia. Postoperative analgesia was provided with acetaminophen administered in the drinking water. Sham-operated animals underwent the same anesthesia, cervical incision and microsurgical exposure of the deep cervical vein and adjacent lymphatic vessel, but no vessel transection or anastomosis was performed. The incision was then closed using the same procedure, and postoperative management was identical to that used for animals in the LVA group.

### Gait analysis

Gait performance was assessed using a rodent gait analysis system (SA114, Sans Biotechnology, China). Measurements were obtained at baseline (before surgery) and 14 days after LVA. Animals were habituated to the testing apparatus for several days before data acquisition to ensure consistent traversal of the walkway. During testing, rats voluntarily crossed a transparent walkway while paw placements were recorded by a high-speed camera positioned beneath the platform. Only uninterrupted runs performed at a steady speed, without pausing or turning, were considered valid for analysis. Multiple valid runs were acquired from each animal and averaged to generate a representative value. Gait data were processed using the manufacturer’s software (Rodent Gait Fine Analysis System) for automated paw detection and parameter extraction. All automated detections were visually reviewed, and manual corrections were applied when necessary to ensure accurate paw assignment. The following gait parameters were quantified: walking speed (cm/s), stride length (cm), mean swing speed (cm/s), stride cycle duration (s), and double support phase for diagonal limb pairs (LF–RH and RF–LH, %). All data were acquired and processed using identical settings across groups. All analyses were conducted by investigators blinded to group allocation.

### Magnetic resonance imaging (MRI) acquisition and analysis

Longitudinal MRI was performed using a 7.0-T small-animal MRI system (Bruker BioSpin GmbH, Ettlingen, Germany) to evaluate structural and perfusion changes after LVA in the AD rat model. MRI examinations were conducted at baseline (before surgery) and at 7 and 14 days after LVA surgery.

Animals were anesthetized with isoflurane (3–4% for induction and 1.5–2.0% for maintenance in medical oxygen) and positioned prone in a dedicated animal holder. Respiration was continuously monitored throughout image acquisition, and body temperature was maintained at 37.0 ± 0.5 °C using a circulating warm-water system.

Structural images were acquired in the axial plane using a T2-weighted TurboRARE sequence to assess brain morphology and ventricular size. Cerebral perfusion was evaluated using two arterial spin labeling sequences (ASL_axial_2seg and pCASL_BBB). Imaging parameters were kept constant across all animals and time points.

MRI data were exported in DICOM format and analyzed using RadiAnt DICOM Viewer (Medixant, Poznań, Poland). Regions of interest were manually delineated by investigators blinded to group allocation. Hippocampal area, lateral ventricle area and third ventricle area were quantified on anatomically matched sections with reference to a standardized rat brain atlas. Bilateral measurements were obtained where applicable and averaged across sections for each animal.

### Immunohistochemistry

Brain tissues of AD rats were collected at predefined time points after LVA surgery, embedded in OCT compound and snap-frozen. Coronal cryosections (20 μm) were prepared and mounted onto adhesive glass slides. Sections were air-dried, permeabilized with 0.1% Triton X-100 and treated with 3% hydrogen peroxide to quench endogenous peroxidase activity. After blocking with 10% normal serum, sections were incubated overnight at 4 °C with primary antibodies (Supplementary Table X), followed by incubation with HRP-conjugated secondary antibodies at 37 °C.

Immunoreactivity was visualized using 3,3′-diaminobenzidine (DAB), and sections were counterstained with hematoxylin, dehydrated and coverslipped. Brightfield images were acquired using an Olympus CX-31 microscope and a NanoZoomer S360 slide scanner. Image acquisition and quantitative analysis were performed using identical settings across groups, with investigators blinded to experimental condition. DAB-positive signal above background was quantified in QuPath (v.0.5.1) as the percentage of positive area.

### Immunofluorescence

At 14 days after LVA surgery, AD rats were deeply anesthetized and transcardially perfused with phosphate-buffered saline (PBS), followed by 4% paraformaldehyde (PFA). Brains were post-fixed in 4% PFA at 4 °C overnight and cryoprotected sequentially in 20% and 30% sucrose in PBS until fully equilibrated. Tissue blocks were embedded in OCT compound and coronally sectioned at 10–15 μm on a cryostat. For antigen retrieval, sections were incubated in Tris–EDTA buffer (pH 9.0) at 95 °C for 15 min and then allowed to cool to room temperature. Sections were blocked for 1 h at room temperature in PBS containing 3% bovine serum albumin (BSA) and 0.3% Triton X-100. Primary antibodies, including anti-transthyretin (TTR) combined with either anti-clusterin (ApoJ/Clu) or anti-vascular endothelial growth factor (VEGF), were applied overnight at 4 °C. After three washes in PBS, sections were incubated with the appropriate fluorophore-conjugated secondary antibodies for 1 h at room temperature. Nuclei were counterstained with DAPI. Fluorescence images were acquired using a Zeiss LSM 980 AiryScan2 confocal microscope with identical acquisition settings across all groups. Multiple sections encompassing anatomically matched regions of the choroid plexus were analyzed for each animal.

### Plasma biomarker measurement by single-molecule detection

Peripheral blood was collected from AD rats in the LVA and sham groups at baseline (before surgery) and at 3, 6, 9 and 12 days after surgery by tail vein puncture. Blood samples were centrifuged at 3,000g for 10 min at 4 °C, and plasma was collected and stored at −80 °C until analysis. Plasma concentrations of phosphorylated tau (pTau181 and pTau217), total tau, Aβ42 and vascular endothelial growth factor (VEGF) were measured using a fully automated single-molecule detection platform (AST-Dx-90, Suzhou AstraBio Technology Co., Ltd.) according to the manufacturer’s instructions. Briefly, 25 μl of plasma was incubated with magnetic beads coated with capture antibodies (Reagent 1) for 6 min, followed by incubation with fluorophore-labeled detection antibodies (Reagent 2) for 4 min at 40 °C. After magnetic separation and washing, fluorescent single-molecule signals were acquired using the integrated imaging module of the instrument. Protein concentrations were calculated from standard curves generated using recombinant protein standards. All samples were processed using identical assay settings, and measurements were performed by investigators blinded to group allocation. For longitudinal analysis, plasma biomarker values were normalized to the corresponding pre-surgical baseline for each animal to reduce inter-individual variability.

### Hydrostatic pressure stimulation, ELISA and bulk RNA sequencing

Z310 cells were maintained in Dulbecco’s modified Eagle medium (DMEM) supplemented with 10% fetal bovine serum at 37 °C in a humidified incubator with 5% CO₂ and grown to 70–80% confluence before treatment. For hydrostatic pressure stimulation, cells were washed with phosphate-buffered saline (PBS) and cultured either in serum-free medium for ELISA experiments or in complete medium for transcriptomic analysis. Cyclic hydrostatic pressure was applied for 24 h using a pressure-loading system at amplitudes of 0.3, 0.5 or 1.0 kPa. For ELISA experiments, stimulation was performed at frequencies of 1.0 or 1.5 Hz; for bulk RNA sequencing, a frequency of 1.5 Hz was used. Control cells were maintained in parallel under static conditions. For secreted protein analysis, culture supernatants were collected after stimulation, centrifuged at 1,000g for 10 min to remove cellular debris and stored at −80 °C until analysis. Clusterin (CLU) and vascular endothelial growth factor (VEGF) concentrations were quantified using commercially available enzyme-linked immunosorbent assay (ELISA) kits according to the manufacturers’ instructions. Absorbance was measured with a microplate reader, and protein concentrations were calculated from standard curves.

For transcriptomic profiling, cells were washed with cold PBS immediately after stimulation and lysed directly in culture plates using TRIzol reagent. Total RNA was extracted according to the manufacturer’s instructions. RNA concentration and purity were assessed using a NanoDrop spectrophotometer, and RNA integrity was evaluated using an Agilent Bioanalyzer. Only samples meeting predefined RNA quality criteria were used for library preparation. Sequencing libraries were generated using a poly(A) enrichment workflow and sequenced on an Illumina platform to obtain paired-end reads. Raw reads were processed to remove adapter sequences and low-quality bases, aligned to the reference genome and quantified at the gene level using a standardized analysis pipeline. Differential gene expression analysis was performed using established workflows with correction for multiple testing. All samples were processed under identical experimental conditions, and all analyses were performed using consistent parameters across groups. Each condition included three independent biological replicates.

### Single-cell RNA sequencing of meninges and hippocampus

Fourteen days after surgery, AD rats from the LVA group (n = 3) and non-surgical control group (n = 3) were euthanized, and the meninges and hippocampi were rapidly dissected on ice and processed immediately for single-cell RNA sequencing. Tissues were transferred into ice-cold calcium- and magnesium-free phosphate-buffered saline (PBS), minced into approximately 0.5-mm fragments and enzymatically dissociated at 37 °C with gentle agitation using a digestion cocktail containing collagenase IV, papain and DNase I. Enzymatic dissociation was terminated with PBS supplemented with fetal bovine serum, followed by gentle trituration to generate single-cell suspensions. Cell suspensions were passed through cell strainers, centrifuged and subjected to red blood cell lysis and dead-cell removal. Cell viability was assessed by trypan blue exclusion, and only samples with viability greater than 85% were used for library preparation. Single-cell suspensions were loaded onto the Chromium Controller (10x Genomics) using the Chromium Single Cell 3′ Reagent Kit v3, with a target recovery of approximately 8,000 cells per sample. cDNA synthesis and library construction were performed according to the manufacturer’s instructions. Libraries were sequenced on an Illumina NovaSeq 6000 platform using paired-end 150-bp reads to a minimum depth of 20,000 reads per cell.

### Bioinformatic analysis

Raw sequencing data were processed using Cell Ranger (v7.0.0, 10x Genomics) for alignment to the reference genome and generation of gene–cell count matrices. Downstream analyses were performed in Seurat (v4.1.0). Low-quality cells were excluded on the basis of predefined quality-control criteria, including fewer than 500 detected genes per cell or greater than 25% mitochondrial gene content. After quality filtering, expression data were normalized and used for principal component analysis, unsupervised clustering and dimensionality reduction by UMAP or t-SNE. Cluster-enriched marker genes were identified and used for cell type annotation.

## Acknowledgments

The authors thank the Science Experiment Center at Hangzhou Institute of Medicine, Chinese Academy of Sciences, for technical support. The authors thank the continued inspiration of Dr. Judah Folkman.

## Author Contributions

X.P., Y.L., L.Y., YJ.Z. and P.G. conceived and designed the study. X.P., S.H., Z.Z., J.Z., Y.H., Z.Z., R.F., X.Y. and Y.L. collected and managed the data. L.Y. performed the statistical analyses. All authors reviewed the manuscript. Y.L., L.Y., YJ.Z. and P.G. organized and supervised this study.

## References

1 Scheltens, P. et al. Alzheimer’s disease. Lancet 397, 1577–1590 (2021). 10.1016/s0140-6736(20)32205-4

2 Zhi, N. et al. The China Alzheimer Report 2025. Gen Psychiatr 38, e102020 (2025). 10.1136/gpsych-2024-102020

3 Ji, Q., Chen, J., Li, Y., Tao, E. & Zhan, Y. Incidence and prevalence of Alzheimer’s disease in China: a systematic review and meta-analysis. Eur J Epidemiol 39, 701–714 (2024). 10.1007/s10654-024-01144-2

4 Jeremic, D., Navarro-López, J. D. & Jiménez-Díaz, L. Efficacy and safety of anti-amyloid-β monoclonal antibodies in current Alzheimer’s disease phase III clinical trials: A systematic review and interactive web app-based meta-analysis. Ageing Res Rev 90, 102012 (2023). 10.1016/j.arr.2023.102012

5 Jucker, M. & Walker, L. C. Alzheimer’s disease: From immunotherapy to immunoprevention. Cell 186, 4260–4270 (2023). 10.1016/j.cell.2023.08.021

6 van Dyck, C. H. et al. Lecanemab in Early Alzheimer’s Disease. N Engl J Med 388, 9–21 (2023). 10.1056/NEJMoa2212948

7 Terao, I. & Kodama, W. Comparative efficacy, tolerability and acceptability of donanemab, lecanemab, aducanumab and lithium on cognitive function in mild cognitive impairment and Alzheimer’s disease: A systematic review and network meta-analysis. Ageing Res Rev 94, 102203 (2024). 10.1016/j.arr.2024.102203

8 Sims, J. R. et al. Donanemab in Early Symptomatic Alzheimer Disease: The TRAILBLAZER-ALZ 2 Randomized Clinical Trial. Jama 330, 512–527 (2023). 10.1001/jama.2023.13239

9 Zhang, J. et al. Recent advances in Alzheimer’s disease: Mechanisms, clinical trials and new drug development strategies. Signal Transduct Target Ther 9, 211 (2024). 10.1038/s41392-024-01911-3

10 Heckmann, B. L. et al. LC3-Associated Endocytosis Facilitates β-Amyloid Clearance and Mitigates Neurodegeneration in Murine Alzheimer’s Disease. Cell 178, 536–551.e514 (2019). 10.1016/j.cell.2019.05.056

11 Fang, E. F. et al. Mitophagy inhibits amyloid-β and tau pathology and reverses cognitive deficits in models of Alzheimer’s disease. Nat Neurosci 22, 401–412 (2019). 10.1038/s41593-018-0332-9

12 Divecha, Y. A. et al. The microcirculation, the blood-brain barrier, and the neurovascular unit in health and Alzheimer disease: The aberrant pericyte is a central player. Pharmacol Rev 77, 100052 (2025). 10.1016/j.pharmr.2025.100052

13 Xie, Q., Pak, C. J., Kwon, J., Chao, S. C. & Hong, J. P. Potential Role of Lymphovenous Bypass in Mitigating Alzheimer’s Disease Dementia. Arch Plast Surg 52, 247–252 (2025). 10.1055/a-2627-9243

14 Yen, Y. H., Chen, M. W., Lim, J. X. & Chew, K. Y. Exploring Lymphovenous Anastomosis for Alzheimer Disease: Addressing Brain Lymphatic Dysfunction, Feasibility, and Outcome Metrics. Plast Reconstr Surg 157, 573–581 (2026). 10.1097/prs.0000000000012364

15 Louveau, A. et al. Structural and functional features of central nervous system lymphatic vessels. Nature 523, 337–341 (2015). 10.1038/nature14432

16 Keil, S. A., Jansson, D., Braun, M. & Iliff, J. J. Glymphatic dysfunction in Alzheimer’s disease: A critical appraisal. Science 389, eadv8269 (2025). 10.1126/science.adv8269

17 Da Mesquita, S., Fu, Z. & Kipnis, J. The Meningeal Lymphatic System: A New Player in Neurophysiology. Neuron 100, 375–388 (2018). 10.1016/j.neuron.2018.09.022

18 Ding, J., Zhao, C., Hao, X. & Jiao, H. Glymphatic and meningeal lymphatic dysfunction in Alzheimer’s disease: Mechanisms and therapeutic perspectives. Alzheimers Dement 21, e70709 (2025). 10.1002/alz.70709

19 Zhao, D. et al. Targeting the glymphatic system: Aβ accumulation and phototherapy strategies across different stages of Alzheimer’s disease. Transl Neurodegener 14, 49 (2025). 10.1186/s40035-025-00510-8

20 Tang, C. et al. Deep cervical lymphatic-venous anastomosis in dementia: a clinical and mechanistic evaluation. Int J Surg 112, 9002–9014 (2025). 10.1097/js9.0000000000004561

21 Fieldhouse, R. China is waging war on Alzheimer’s. What can its approach teach the rest of the world? Nature 650, 816–818 (2026). 10.1038/d41586-026-00564-2

22 Da Mesquita, S. et al. Functional aspects of meningeal lymphatics in ageing and Alzheimer’s disease. Nature 560, 185–191 (2018). 10.1038/s41586-018-0368-8

23 Zhou, J. et al. BACE1 regulates expression of Clusterin in astrocytes for enhancing clearance of β-amyloid peptides. Mol Neurodegener 18, 31 (2023). 10.1186/s13024-023-00611-w

24 Liu, H. et al. APOE3ch alleviates Aβ and tau pathology and neurodegeneration in the human APP(NL-G-F) cerebral organoid model of Alzheimer’s disease. Cell Res 34, 451–454 (2024). 10.1038/s41422-024-00957-w

25 Feng, S. et al. High-intensity interval training ameliorates Alzheimer’s disease-like pathology by regulating astrocyte phenotype-associated AQP4 polarization. Theranostics 13, 3434–3450 (2023). 10.7150/thno.81951

26 Ma, Y. et al. Targeting blood brain barrier-Remote ischemic conditioning alleviates cognitive impairment in female APP/PS1 rats. CNS Neurosci Ther 30, e14613 (2024). 10.1111/cns.14613

27 Fu, X. et al. Deep cervical lymphatic-venous anastomosis attenuates cognitive dysfunction and biomarker abnormalities in severe Alzheimer’s disease: A prospective single-arm study. Alzheimers Dement 22, e71150 (2026). 10.1002/alz.71150

28 Kim, M. et al. Surface-functionalized SERS platform for deep learning-assisted diagnosis of Alzheimer’s disease. Biosens Bioelectron 251, 116128 (2024). 10.1016/j.bios.2024.116128

29 Youmans, K. L. et al. Intraneuronal Aβ detection in 5xFAD mice by a new Aβ-specific antibody. Mol Neurodegener 7, 8 (2012). 10.1186/1750-1326-7-8

30 Ramesha, S. et al. Unique molecular characteristics and microglial origin of Kv1.3 channel-positive brain myeloid cells in Alzheimer’s disease. Proc Natl Acad Sci U S A 118 (2021). 10.1073/pnas.2013545118

31 Tian, C. et al. Understanding monocyte-driven neuroinflammation in Alzheimer’s disease using human cortical organoid microphysiological systems. Sci Adv 11, eadu2708 (2025). 10.1126/sciadv.adu2708

32 Chen, J. Y. et al. Deep cervical lymphovenous anastomosis (LVA) for Alzheimer’s disease: microsurgical procedure in a prospective cohort study. Int J Surg 111, 4211–4221 (2025). 10.1097/js9.0000000000002490

33 Lau, S. F., Cao, H., Fu, A. K. Y. & Ip, N. Y. Single-nucleus transcriptome analysis reveals dysregulation of angiogenic endothelial cells and neuroprotective glia in Alzheimer’s disease. Proc Natl Acad Sci U S A 117, 25800–25809 (2020). 10.1073/pnas.2008762117

34 Serrano-Pozo, A., Das, S. & Hyman, B. T. APOE and Alzheimer’s disease: advances in genetics, pathophysiology, and therapeutic approaches. Lancet Neurol 20, 68–80 (2021). 10.1016/s1474-4422(20)30412-9

35 Long, J. M. & Holtzman, D. M. Alzheimer Disease: An Update on Pathobiology and Treatment Strategies. Cell 179, 312–339 (2019). 10.1016/j.cell.2019.09.001

36 Morabito, S. et al. Single-nucleus chromatin accessibility and transcriptomic characterization of Alzheimer’s disease. Nat Genet 53, 1143–1155 (2021). 10.1038/s41588-021-00894-z

37 Desikan, R. S. et al. The role of clusterin in amyloid-β-associated neurodegeneration. JAMA Neurol 71, 180–187 (2014). 10.1001/jamaneurol.2013.4560

38 Zeng, X. et al. Multi-analyte proteomic analysis identifies blood-based neuroinflammation, cerebrovascular and synaptic biomarkers in preclinical Alzheimer’s disease. Mol Neurodegener 19, 68 (2024). 10.1186/s13024-024-00753-5

39 Monroe, K. M., Hong, S., Lewcock, J. W. & Yang, A. C. Therapeutic targeting of neuroimmune mechanisms in neurodegeneration. Nat Rev Drug Discov 25, 390–405 (2026). 10.1038/s41573-025-01370-7

40 Murdock, M. H. et al. Multisensory gamma stimulation promotes glymphatic clearance of amyloid. Nature 627, 149–156 (2024). 10.1038/s41586-024-07132-6

41 Boespflug, E. L. & Iliff, J. J. The Emerging Relationship Between Interstitial Fluid-Cerebrospinal Fluid Exchange, Amyloid-β, and Sleep. Biol Psychiatry 83, 328–336 (2018). 10.1016/j.biopsych.2017.11.031

42 Nedergaard, M. & Goldman, S. A. Glymphatic failure as a final common pathway to dementia. Science 370, 50–56 (2020). 10.1126/science.abb8739

43 Jiang-Xie, L. F. et al. Neuronal dynamics direct cerebrospinal fluid perfusion and brain clearance. Nature 627, 157–164 (2024). 10.1038/s41586-024-07108-6

44 Ding, X. B. et al. Impaired meningeal lymphatic drainage in patients with idiopathic Parkinson’s disease. Nat Med 27, 411–418 (2021). 10.1038/s41591-020-01198-1

45 Gao, F. et al. Pathological angiogenesis was associated with cerebrovascular lesion and neurodegeneration in Alzheimer’s disease. Alzheimers Dement 21, e14521 (2025). 10.1002/alz.14521

46 Taghvaei, M. et al. Regional cerebral blood flow reflects both neurodegeneration and microvascular integrity across the Alzheimer’s continuum. Alzheimers Dement 21, e14382 (2025). 10.1002/alz.14382

47 Robert, S. M. et al. The choroid plexus links innate immunity to CSF dysregulation in hydrocephalus. Cell 186, 764–785.e721 (2023). 10.1016/j.cell.2023.01.017

48 Courtney, Y. et al. Choroid plexus apocrine secretion shapes CSF proteome during mouse brain development. Nat Neurosci 28, 1446–1459 (2025). 10.1038/s41593-025-01972-9

49 Yeh, F. L., Wang, Y., Tom, I., Gonzalez, L. C. & Sheng, M. TREM2 Binds to Apolipoproteins, Including APOE and CLU/APOJ, and Thereby Facilitates Uptake of Amyloid-Beta by Microglia. Neuron 91, 328–340 (2016). 10.1016/j.neuron.2016.06.015

50 Lee, C. et al. Vascular endothelial growth factor signaling in health and disease: from molecular mechanisms to therapeutic perspectives. Signal Transduct Target Ther 10, 170 (2025). 10.1038/s41392-025-02249-0

51 Martins, L. F. et al. Motor neurons use push-pull signals to direct vascular remodeling critical for their connectivity. Neuron 110, 4090–4107.e4011 (2022). 10.1016/j.neuron.2022.09.021

52 Zhu, X. et al. Piezo1-mediated mechanotransduction in choroid plexus epithelial cells governs ciliogenesis and cerebrospinal fluid homeostasis. Neuron (2026). 10.1016/j.neuron.2026.02.033

